# Molecular signature of COVID-19 prior to its exacerbation by multi-omics survey

**DOI:** 10.64898/2025.12.27.696524

**Authors:** Tomoyuki Suzuki, Yoshihiro Kita, Keisuke Yanagida, Kenji Maeda, Tomomi Hashidate-Yoshida, Hiroki Nakanishi, Takayo Ohto-Nakanishi, Junko Terada-Hirashima, Yoshie Tsujimoto, Masayuki Hojo, Hiroaki Mitsuya, Takao Shimizu, Hideo Shindou

## Abstract

Severe acute respiratory syndrome coronavirus 2 (SARS-CoV-2) caused a global pandemic due to its high transmissibility and ability to evade innate immune responses. Comprehensive characterization of the disease is essential for elucidating its pathophysiology and clinical progression. In this study, we performed multi-omics analyses of plasma samples collected from SARS-CoV-2-positive patients prior to clinical deterioration of coronavirus disease 2019 (COVID-19). These samples revealed the potential of previously reported clinical parameters, including CRP and neutrophil level, to predict COVID-19 exacerbation in the early stage. Our analysis identified a novel panel of molecules that precede the clinical manifestations associated with COVID-19 progression. These candidate biomarkers exhibited strong correlations with previously reported clinical and immunological parameters. Notably, several inflammation-related markers showed inverse associations with specific interferon subtypes, including IFN-α6 and IFN-α8, potentially reflecting mechanisms of SARS-CoV-2-mediated immune evasion. Our findings contribute to the understanding of virus-induced acute exacerbation and offer a valuable foundation for future pandemic research.

## 1. Introduction

Severe acute respiratory syndrome coronavirus 2 (SARS-CoV-2), the causative agent of coronavirus disease 2019 (COVID-19), has affected billions of people worldwide. Patients with COVID-19 exhibited higher in-hospital mortality rates (∼1–2%) compared to those with influenza virus infections [1–3]. Numerous studies have identified advanced age, male sex, and certain comorbidities as risk factors for severe COVID-19 [4,5]. In response, the development and widespread use of highly effective vaccines and newly introduced oral antiviral agents have significantly reduced hospitalization rates and mortality associated with COVID-19 [6,7]. Although the global health emergency status of COVID-19 was declared over in May 2023, several challenges remain, including long COVID and the recurrent emergence of highly transmissible SARS-CoV-2 variants such as JN.1 (BA.2.86.1.1) and NB.1.8.1 (a descendant of BA.2.86) [8].

Two major factors contributed to the global spread and impact of COVID-19: its high transmissibility during the presymptomatic phase and its potential for sudden clinical deterioration. During the early stages of the pandemic, extensive research focused on identifying high-risk individuals and effective treatment strategies. These efforts revealed that innate immune deficiencies and cytokine release syndrome (CRS) are key features of severe COVID-19 pathology [9–12]. In particular, deficiencies in type I and III interferon (IFN) responses have been strongly associated with increased susceptibility to severe SARS-CoV-2 infection [10,13]. SARS-CoV-2 is known to subvert host immune defenses by inhibiting mRNA splicing, translation, and nuclear translocation of transcription factors [14–16]. CRS, another hallmark of severe COVID-19, represents a hyperinflammatory state characterized by excessive production of cytokines. Elevated levels of proinflammatory cytokines such as interleukin (IL)-6, interferon-γ (IFN-γ), and tumor necrosis factor-α (TNF-α) have been correlated with disease severity in numerous studies [12,17–19].

To date, multi-omics approaches—including proteomics, metabolomics, and lipidomics—have uncovered molecular signatures associated with COVID-19 pathogenesis and identified candidate biomarkers for disease severity [20–23]. Longitudinal multi-omics studies have further highlighted important pathological features, including epithelial damage and disrupted lipid transport [24,25]. In this study, we focused on the early stage of COVID-19 progression to identify molecular signature of its exacerbation and find novel biomarkers for further COVID-19 studies.

## 2. Materials and Methods

### 2.1. Human subjects and sample collection

Between January and December 2020, patients diagnosed with SARS-CoV-2 infection by polymerase chain reaction (PCR) testing were admitted to the National Center for Global Health and Medicine (NCGM), Tokyo, Japan. A total of 399 individuals were enrolled in a prospective observational study (NCGM-G-003472-03), and written informed consent was obtained from all participants. Based on the plasma samples collected during this prospective cohort, we conducted a retrospective analysis to investigate the molecular signatures associated with COVID-19. The study protocol was approved by the institutional ethics committee (NCGM-G-004061-02). The name of our institute has been changed from NCGM to Japan Institute for Health Security.

Participants were eligible for inclusion if they were hospitalized with mild or moderate COVID-19 symptoms and had plasma samples collected on the first day of admission. Among the 399 Japanese patients enrolled in the prospective observational study, 7 individuals were classified as “progressive” cases. These individuals initially presented with mild or moderate symptoms but later developed severe or critical illness during hospitalization. Notably, all patients in the progressive group were male. Considering that both age and biological sex are known risk factors for severe COVID-19, a control group of 23 “stable” patients matched for age and sex was selected. These control patients remained clinically stable throughout hospitalization and were discharged without progressing to severe or critical disease. Disease severity was categorized into four stages—mild, moderate, severe, and critical—based on the National Institutes of Health (NIH) guidelines (https://www.covid19treatmentguidelines.nih.gov/). Briefly, mild cases included patients with COVID-19-related symptoms but without dyspnea, shortness of breath, or abnormal chest imaging. Moderate illness was defined as evidence of lower respiratory tract disease with a peripheral oxygen saturation (SpO₂) ≥94% on room air. Severe cases exhibited at least one of the following: SpO₂ <94% on room air, PaO₂/FiO₂ ratio <300 mmHg, respiratory rate >30 breaths/min, or lung infiltrates involving >50% of the lungs. Critical illness was defined by the presence of respiratory failure, septic shock, and/or multiple organ dysfunction. EDTA-anticoagulated plasma samples were collected on the day of admission and at subsequent time points during hospitalization. All plasma samples were aliquoted and stored at –80 °C until further analysis. COVID-19 REGISTRY JAPAN (COVIREGI-JP) data of National Center for Global Health and Medicine were used for this study with permission.

### 2.2 Lipidomics sample preparation and measurement of lipids

A 10 µl aliquot was diluted 10-fold with Buffer AVL (QIAGEN, Cat#19089), and then 1.5 ml MeOH which included 31 internal standards was added. Comprehensive lipids were conducted as previously described [26]. Briefly, total lipids were extracted with the Bligh-Dyer method [27]. An aliquot of the lower/organic phase was evaporated to dryness under N2, and the residue was dissolved in methanol for liquid chromatography-mass spectrometry (LC-MS) measurements of phosphatidylcholine (PC) and phosphatidylethanolamine (PE). To analyze phosphatidic acid (PA), phosphatidylserine (PS), phosphatidylglycerol (PG) and phosphatidylinositol (PI), another aliquot of the same lipid extract was added with an equal volume of the methanol before being loaded onto a DEAE-cellulose column (Santa Cruz Biotechnology) pre-equilibrated with chloroform. After successive washes with chloroform/methanol (1:1, v/v), the acidic phospholipids (PLs) were eluted with chloroform/methanol/HCl/water (12:12:1:1, v/v), followed by evaporation to dryness to give a residue, which was redissolved in methanol.

LC-electrospray ionization-MS/MS analysis was performed consisting of ACQUITY UPLC H-Class (Waters) and a triple quadrupole mass spectrometer Xeno TQ-XS (Waters). The lipid samples were separated on an X-Bridge C18 column (Waters, Cat#186003128) at 40 °C using a gradient solvent system as follows: mobile phase A (isopropyl alcohol/methanol/water (5:1:4 v/v/v) supplemented with 5 mM ammonium formate and 0.05% ammonium hydroxide) and mobile phase B (isopropyl alcohol supplemented with 5 mM ammonium formate and 0.05% ammonium hydroxide) ratios of 70:30 (0 min), 50:50 (2 min), 20:80 (13 min), 5:95 (15-30 min), 95:5 (31-35 min), and 70:30 (35-45 min). The flow rate was 0.08 ml/min. Peak areas of individual species were normalized against a sum of the detected signals for each sample. Raw data were analyzed by using MassLynx4.2 (Waters).

### 2.3 Lipidomics sample preparation and measurement of lipid mediators

A 160 µl aliquot was diluted 10-fold with Buffer AVL (QIAGEN, Cat#19089), and then was further diluted 5-fold by MeOH with a deuterium-labeled internal standard mixture, which included PAF-d4, LysoPAF-d4, PGA2-d4, PGD2-d4, PGE2-d4, PGF2α-d9, 6-keto-PGF1α-d4, TXB2-d4, LTB4-d4, LTC4-d5, LTD4-d5, LTE4-d5, 5-HETE-d8, 12-HETE-d8, 15-HETE-d8, AEA-d4, OEA-d4, tetranor-PGEM-d6, AA-d8, EPA-d5, DHA-d5, 14,15-DiHET-d11 and 11,12-EET-d11. The supernatants were collected after centrifugation at 20,000 g for 15 min at 4 °C. The extracts were purified by a solid-phase method using an Oasis HLB column (Waters, Cat#186000381).

Lipid mediators were measured using a triple quadrupole mass spectrometer LCMS-8060 (Shimadzu) as previously described [28,29]. Briefly, a reverse-phase column (Phenomenex, Cat#00F-4497-AN) was used for chromatographic separation with a binary mobile phase comprising 0.1% formic acid/water (mobile phase A) and acetonitrile (mobile phase B). The gradient of the mobile phase (%A/%B) was programmed as follows: 0 min (90/10), 5 min (75/25), 10 min (65/35), 20 min (25/75), 20-28 min (5/95), 28-30 min (90/10).

The flow rate was 0.4 ml/min, and the column temperature was 40 ℃. Raw data were analyzed by using LabSolutions Insight (Shimadzu). The signals were compared with standard curves for quantification.

### 2.4 Metabolomics sample preparation and measurement

A 50 µl aliquot was diluted 5-fold methanol which included internal standards (Human Metabolome Technologies, Inc. (HMT)). 150 µl Milli-Q water was added before the sample was centrifuged at 9,100 g with an ultrafiltration filter to remove proteins. The filtrate was centrifugally concentrated and re-suspended in 50 µl of Milli-Q water for capillary electrophoresis time-of-flight mass spectrometry (CE-TOFMS) analysis.

Metabolome analysis was conducted by the Basic Scan package from HMT using CE-TOFMS based on previously described methods [30,31]. Briefly, CE-TOFMS analysis was conducted using an Agilent CE capillary electrophoresis system equipped with an Agilent 6210 time-of-flight mass spectrometer (Agilent Technologies, Inc.). The spectrometer was scanned from 50 to 1,000 m/z, and peaks were extracted using MasterHands ver.2.18.0.1 automatic integration software (Keio University) to obtain peak information including m/z, peak area, and migration time (MT) [32]. Signal peaks corresponding to isotopomers, adduct ions and other product ions of known metabolites were excluded, and based on their m/z values with the MTs, remaining peaks were annotated according to the HMT’s proprietary metabolite database. The areas of the annotated peaks were normalized based on internal standard levels and sample quantities to obtain relative levels of each metabolite.

### 2.5 Proteomics sample preparation and measurement

130 µL EDTA plasma samples from 30 participants were sent to SomaLogic and processed using the SomaScan 7k Assay (v4.1) [33]. SomaScan uses single-stranded DNA-based aptamers called Slow Off-rate Modified Aptamers (SOMAmer reagents) to simultaneously measure thousands of human proteins ranging from femto- to micro-molar concentrations. The modified aptamer binding reagents [34], SomaScan assay [35], its performance characteristics [33,36], and specificity [37–39] to human targets have been previously described. The assay uses standard controls including 12 hybridization normalization control sequences used to control for variability in the Agilent readout process as well as 5 human calibrator control pooled replicates and 3 quality control pooled replicates used to mitigate batch effects and verify the quality of the assay run using standard acceptance criteria. The readout is performed using Agilent hybridization, scan, and feature extraction technology. The initial raw signal intensity from the Agilent scan is standardized using a series of linear scaling procedures developed to mitigate systematic biases that may lead to excess variation and skewness in the data. Results are reported as Relative Fluorescence Units (RFUs) as a relative measurement of protein abundance.

To avoid bias, laboratory personnel were blinded to sample identity, and samples were randomly arranged on the plates, with a set of calibration and normalization samples. Data from all samples passed quality-control criteria and were fit for analysis.

### 2.6 Data analysis

All statistical analyses were performed using R (version 4.1.2, https://www.r-project.org/). Unless otherwise specified, data are presented as mean ± standard error of the mean (SEM). Statistical significance was defined as *p* < 0.05.

#### 2.6.1 Data preprocessing and feature selection

● Clinical and omics data (lipidomics, metabolomics, proteomics, and lipid mediators) were curated to exclude features with >20% missing values in either group.
● Features with ≤20% missing values were imputed by replacing the missing values with half of the minimal value observed for that feature within the respective group.

#### 2.6.2 Group comparisons and feature extraction

● To identify differentially abundant features between groups (“stable” vs. “progressive”), unpaired two-tailed Student’s t-tests with Welch’s correction were conducted.
● A molecule was considered significantly altered if it met the thresholds of |log2 fold change(FC)| > 0.585 (≥1.5-FC) and *p* < 0.05.
● For multiple comparisons among clinical blood parameters, *p* values were adjusted using the Benjamini–Hochberg (BH) method to control the false discovery rate (FDR).

#### 2.6.3 Regression and correlation analyses

● Univariate logistic regression models with age as a covariate were constructed for each variable to evaluate associations “progressive” COVID-19.
● Variables with odds ratios whose 95% confidence intervals did not include 1 were considered statistically significant and visualized using forest plots.
● Pearson correlation analyses were employed to assess relationships between omics features and clinical parameters.

#### 2.6.4 Multivariate and enrichment analyses

● Partial Least Squares Discriminant Analysis (PLS-DA) was performed using the ropls R package [40].
● Hierarchical clustering was conducted using Ward’s linkage algorithm with Euclidean distance for similarity metrics.
● Metascape (https://metascape.org/) was utilized for functional enrichment analysis of selected proteins [41].

#### 2.6.5 Receiver Operating Characteristic (ROC) curve analysis

● Univariate ROC curve analysis was performed to evaluate the predictive ability of individual variables.
● AUC (Area Under the Curve) values were calculated and visualized using the plotROC R package [42].

## 3. Results

### 3.1 Study design

The goal of this study was to identify molecular features associated with early-stage exacerbation of COVID-19 by analyzing plasma samples collected from SARS-CoV-2-positive patients at the time of hospital admission, prior to the development of severe disease. We adopted NIH guidelines to define severe cases who had SpO₂ < 94% on room air and required supplemental oxygen. To investigate molecular alterations that reflect the initial phase of disease progression, we stratified patients into two groups: (1) the “stable” group, comprising individuals who remained mild or moderate condition throughout hospitalization, and (2) the “progressive” group, consisting of patients who were initially admitted in a mild or moderate state but later progressed to severe disease (Fig. 1.).

**Fig. 1.**
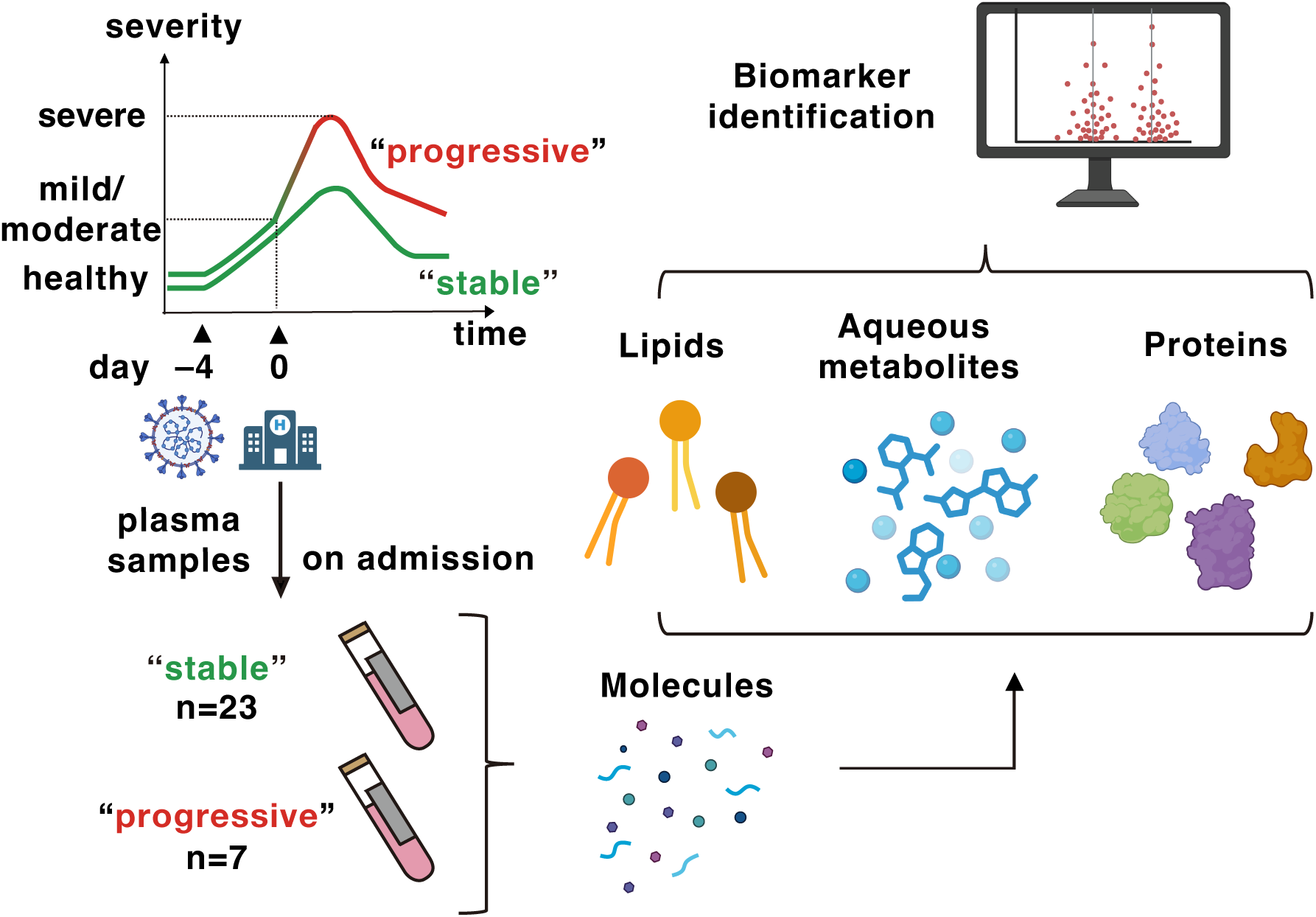
Study design and patient grouping. Multi-omics analyses, including lipidomics, metabolomics, and proteomics, were conducted using plasma samples collected from COVID-19 patients at the time of hospital admission to evaluate a molecular profile of its exacerbation. Of the 30 patients, 23 remained in mild or moderate condition throughout hospitalization and were categorized as the “stable” group. The remaining 7 patients, initially classified as mild or moderate on admission, later progressed to severe or critical illness and were defined as the “progressive” group. The green line indicates mild/moderate clinical status, while the magenta line indicates severe/critical clinical status. Created in BioRender. Suzuki, T. (2025) https://BioRender.com/8x0lfcn

A total of 399 COVID-19 patients were enrolled in a prospective observational study at our hospital between January and December 2020. Among these, 7 patients met the criteria for the “progressive” group, all of whom were male. A control group of 23 age- and sex-matched patients was selected for the “stable” group.

### 3.2 Patient characteristics

Due to the small number of patients in the “progressive” group, no apparent distributions were observed between the groups in terms of smoking history, self-reported medical history, COVID-19-related symptoms, or the time from symptom onset to hospital admission (Table 1). Both groups presented with comparable SpO₂ levels at admission; however, patients in the “progressive” group required oxygen supplementation an average of 4.1 days after admission and had longer hospital stays. None of the “progressive” patients required mechanical ventilation. While only symptomatic treatment was administered to the “stable” group, patients in the “progressive” group received steroids (n = 5, 71.4%), remdesivir (n = 4, 57.1%), or tocilizumab (n = 1, 14.3%). All participants in both groups were discharged without the need for supplemental oxygen.

**Table 1.** Baseline demographic and clinical characteristics of patients. Percentages are calculated within each group. Data are presented as mean ± SD. Abbreviations: N/A, not applicable; BMI, body mass index.

| **Parameter** | **stable** | **progressive** |
| --- | --- | --- |
| Age | 54.8 ± 16.9 | 52.1 ± 5.6 |
| Height (cm) | 170.4 ± 8.1 | 172.5 ± 7.1 |
| Weight (kg) | 73 ± 16.5 | 70.8 ± 12.3 |
| BMI (kg/m<sup>2</sup>) | 25 ± 4.9 | 23.6 ± 2.6 |
| SpO<sub>2</sub> on admission (%) | 97 ± 1.3 | 96.6 ± 1.6 |
| **Social history** |  |  |
| None smoker | 8 (34.8%) | 1 (14.3%) |
| Former smoker | 9 (39.1%) | 3 (42.9%) |
| Current smoker | 6 (26.1%) | 3 (42.9%) |
| **Past medical history** |  |  |
| Heart disease | 3 (13%) | 0 (0%) |
| Lung disease | 6 (26.1%) | 2 (28.6%) |
| Cerebrovascular disease | 1 (4.3%) | 0 (0%) |
| Hypertension | 6 (26.1%) | 1 (14.3%) |
| Diabetes mellitus | 8 (34.8%) | 1 (14.3%) |
| Autoimmune disease | 2 (8.7%) | 0 (0%) |
| Cancer | 1 (4.3%) | 0 (0%) |
| **COVID-19 related symptom** |  |  |
| Fever | 16 (69.6%) | 7 (100%) |
| Cough | 13 (56.5%) | 4 (57.1%) |
| Difficulty in breathing | 4 (17.4%) | 4 (57.1%) |
| Diarrhea | 5 (21.7%) | 2 (28.6%) |
| Taste smell disorder | 4 (17.4%) | 2 (28.6%) |
| Lung Infiltration | 19 (82.6%) | 7 (100%) |
| **Time course** |  |  |
| Onset - Admission (days) | 4.8 ± 3.5 | 3.9 ± 3.1 |
| Admission - Oxygen (days) | N/A | 4.1 ± 2.3 |
| Oxygen duration (days) | N/A | 5 ± 4.5 |
| Length of stay (days) | 10.5 ± 3.9 | 16.1 ± 3.3 |

### 3.3 Laboratory data on admission

Although all patients were classified as having mild to moderate disease on admission, the “progressive” group already showed a reduction in lymphocyte percentage and an increase in neutrophil percentage (Table 2). Inflammatory markers such as C-reactive protein (CRP) and tissue damage marker lactate dehydrogenase (LDH) were also significantly elevated in the “progressive” group. In contrast, liver enzymes (AST, ALT) and the renal function marker creatinine showed no significant differences between the groups.

**Table 2.** Laboratory parameters on admission. Data are presented as mean ± SD.*q*-values for numeric variables were calculated using two-tailed Student’s *t*-test with unequal variance and adjusted for multiple comparisons using the Benjamini–Hochberg method. Significance thresholds: \*\**q* < 0.01, \**q* < 0.1.

|  | stable | progressive | *q value* |
| --- | --- | --- | --- |
| WBC ($10^3/\mu\text{l}$) | $4.4 \pm 1.1$ | $5.7 \pm 1.6$ | 0.28 |
| Neutrophil percentage (%) | $59.7 \pm 8.6$ | $75.6 \pm 7$ | 0.0003** |
| Lymphocyte percentage (%) | $29.2 \pm 8.4$ | $17.7 \pm 5$ | 0.0003** |
| Neutrophil number ($10^3/\mu\text{l}$) | $2.7 \pm 1$ | $4.3 \pm 1.4$ | 0.097* |
| Lymphocyte number ($10^3/\mu\text{l}$) | $1.3 \pm 0.4$ | $1 \pm 0.3$ | 0.172 |
| Hb (g/dL) | $14.8 \pm 1.6$ | $14.6 \pm 1.2$ | 0.98 |
| Ht (%) | $43.3 \pm 4.4$ | $42.9 \pm 2.9$ | 0.98 |
| Plt ($10^3/\mu\text{l}$) | $85.5 \pm 109$ | $86.6 \pm 94.2$ | 0.98 |
| Na (mEq/L) | $139.7 \pm 3$ | $138.1 \pm 3$ | 0.34 |
| K (mEq/L) | $3.9 \pm 0.4$ | $4.2 \pm 0.3$ | 0.45 |
| Albmin (g/dL) | $4 \pm 0.3$ | $3.7 \pm 0.5$ | 0.34 |
| T.Bil (mg/dL) | $0.6 \pm 0.2$ | $0.6 \pm 0.2$ | 0.98 |
| AST (U/L) | $32.7 \pm 14.4$ | $44.6 \pm 30.4$ | 0.56 |
| ALT (U/L) | $41.8 \pm 27.7$ | $42.4 \pm 40$ | 0.98 |
| LDH (U/L) | $198.7 \pm 40.3$ | $287.1 \pm 76.6$ | 0.097* |
| BUN (mg/dL) | $14.2 \pm 4.5$ | $11.8 \pm 3.3$ | 0.34 |
| Cre (mg/dL) | $0.9 \pm 0.2$ | $0.9 \pm 0.1$ | 0.98 |
| Glu (mg/dL) | $114.1 \pm 33.2$ | $128 \pm 58.9$ | 0.9 |
| CRP (mg/dL) | $1.4 \pm 2.8$ | $6.5 \pm 4.2$ | 0.097* |

### 3.4 Altered molecules in the early stage of COVID-19 exacerbation

To investigate molecular changes preceding severe disease, we performed multi-omics analyses—including lipidomics, metabolomics, and proteomics—on plasma samples collected at admission. Lipidomics was further divided into phospholipid and lipid mediator analyses due to differences in molecular polarity and extraction protocols. In PLS-DA, proteomic data clearly distinguished the “progressive” group from the “stable” group, while lipidomic and metabolomic data exhibited less pronounced separation (Fig. 2A.).

**Fig. 2.**
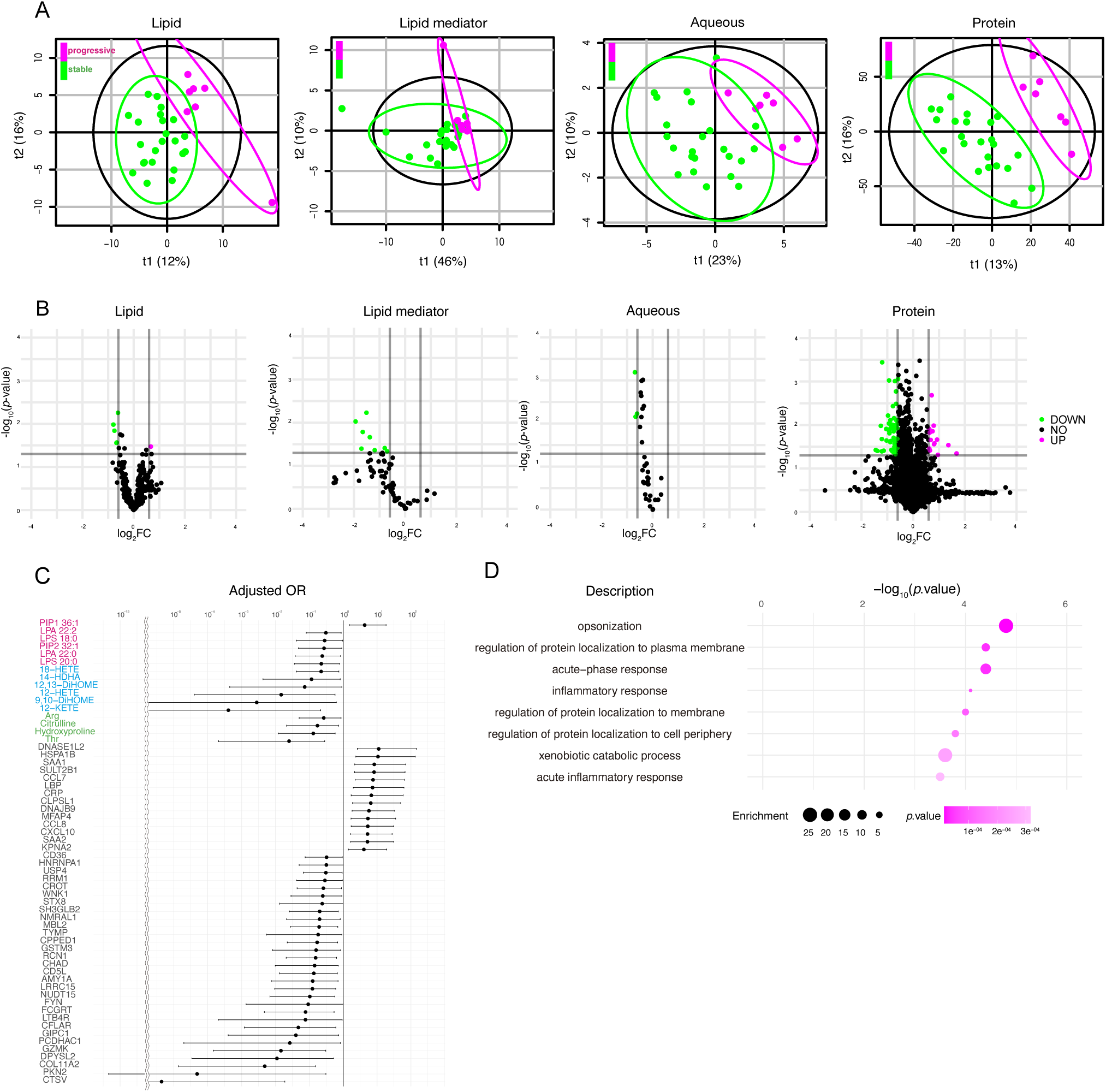
Dysregulated molecules prior to COVID-19 progression. (A) Partial least squares discriminant analysis (PLS-DA) of plasma lipids, lipid mediators, aqueous metabolites, and proteins in the “stable” (green) and “progressive” (magenta) groups. (B) Volcano plots showing differentially abundant lipids, lipid mediators, aqueous metabolites, and proteins between groups (|log2 fold change(FC)| > 0.585 (≥1.5-FC) and *p* < 0.05). (C) Logistic regression analyses adjusted for age were performed for each molecule. Molecules with odds ratios (ORs) whose 95% confidence intervals excluded 1 were shown in forest plots. Lipids are indicated in red, lipid mediators in blue, aqueous metabolites in green, and proteins in black. (D) Gene Ontology enrichment analysis of dysregulated proteins identified in panel (B).

Volcano plots highlighted differentially expressed molecules in each omics layer (Fig. 2B; Tables S1–S4). To refine the list of candidate biomarkers, logistic regression analysis was conducted to calculate adjusted odds ratios, yielding 6 phospholipids, 6 lipid mediators, 4 polar metabolites, and 44 proteins, which were visualized in a forest plot (Fig. 2C). Gene ontology enrichment analysis of the selected proteins revealed the top three enriched biological processes: opsonization, regulation of protein localization to the plasma membrane, and the acute-phase response (Fig. 2D.).

### 3.5 Altered molecules correlated with COVID-19 exacerbation

We next evaluated the potential of these molecules to predict disease progression using receiver operating characteristic (ROC) curve analysis. Molecules with an area under the curve (AUC) > 0.85 included threonine (Thr), C-C motif chemokine ligand 7 (CCL7), CCL8, dnaJ heat shock protein family member B9 (DNAJB9), heat shock protein family A member 1B (HSPA1B), Microfibril associated protein 4 (MFAP4), collagen type XI alpha 2 chain (COL11A2), cathepsin V (CTSV), and deoxyribonuclease 1 like 2 (DNASE1L2) (Fig. 3A–I.). Among them, Thr showed the highest AUC (0.913) and was decreased in the “progressive” group (Fig. 3A.).

**Fig. 3.**
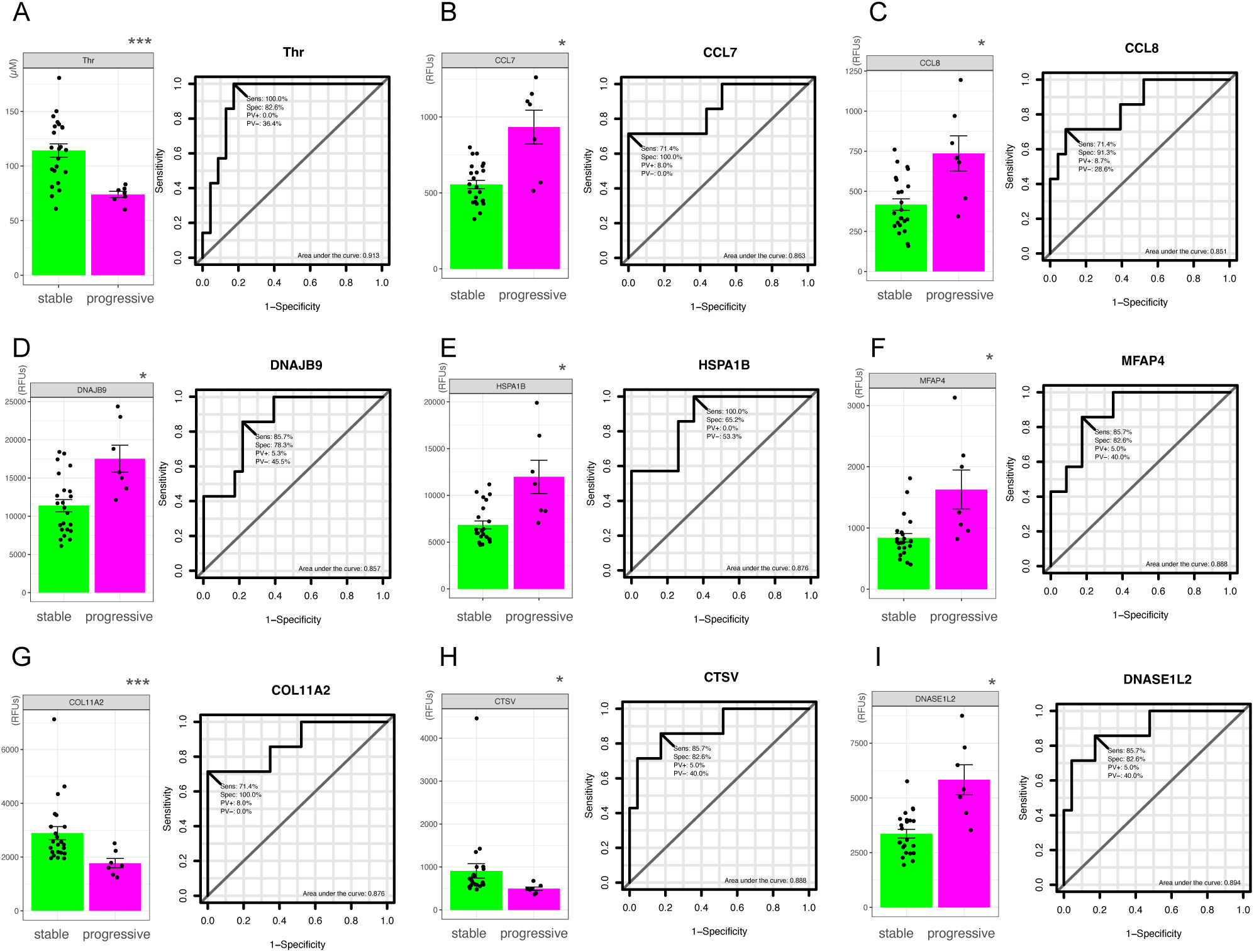
Molecular biomarker candidates for COVID-19 progression. (A–I) Bar plots showing the levels of selected molecules in plasma samples collected on admission (mean ± SEM). Statistical significance is indicated (\*\*\**p* < 0.001, \**p* < 0.05). Corresponding ROC curves are shown for each candidate biomarker.

Logistic regression analysis was also conducted to calculate adjusted odds ratios using clinical parameters (Table S5). Among them, CRP, neutrophil percentage, and lymphocyte percentage also exhibited AUC values > 0.85 (Fig. 4A–C.).

**Fig. 4.**
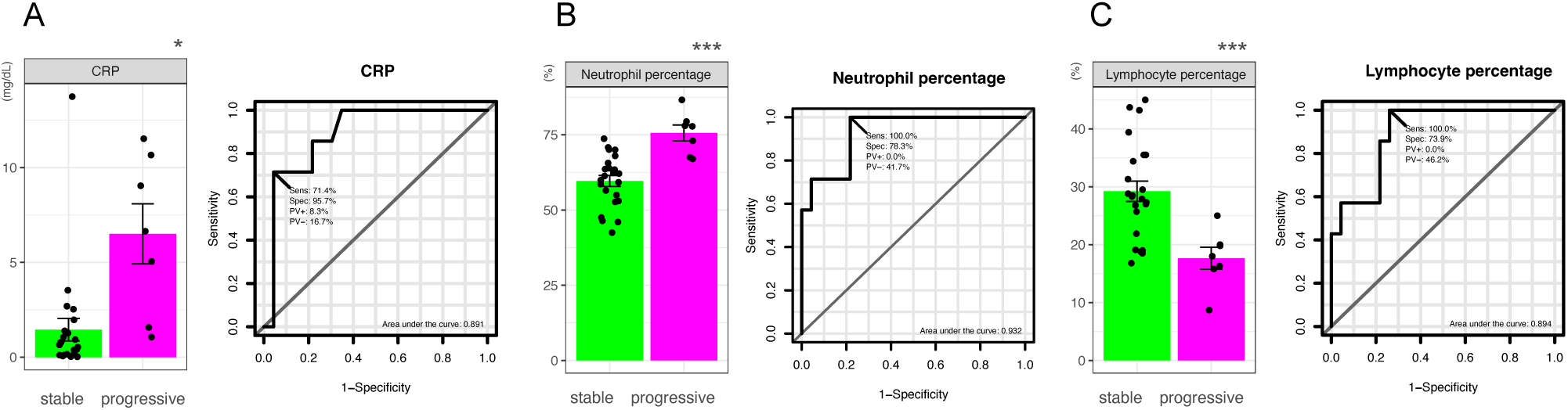
Clinical biomarker candidates for COVID-19 progression. (A–C) Bar plots showing clinical laboratory values on admission (mean ± SEM), along with their ROC curves. Statistical significance is indicated (\*\*\**p* < 0.001, \**p* < 0.05).

In our study, plasma samples were also collected not only on admissions but also around day 7 post-admission. The follow-up measurements after admission revealed that Thr as well as MFAP4 levels did not significantly differ between groups (Fig. S1.), suggesting their utility as early, but not late, biomarkers. Interestingly, CRP levels in the “progressive” group normalized by day 7, diminishing its predictive value at later stages (Fig. S2.).

### 3.6 Associations between altered molecules and clinical/immunological features

CRP, neutrophil percentage, and lymphocyte percentage are well-known markers of COVID-19 severity [43], and the changes in them are unusual for typical viral infections. To explore their relationship with the altered molecules, we computed Pearson correlations between these three parameters and the molecules identified in Fig. 2C (Fig. 5A.; Table S6). Hierarchical clustering revealed distinct patterns. CRP and neutrophil percentage were strongly positively correlated with acute inflammatory proteins, such as CCL7 and serum amyloid A2 (SAA2). Notably, CRP had the strongest correlation with SAA2 (r = 0.93), while neutrophil percentage correlated most strongly with DNAJB9 (r = – 0.68). Conversely, DNAJB4 had the strongest negative correlation with lymphocyte percentage (r = – 0.65).

**Fig. 5.**
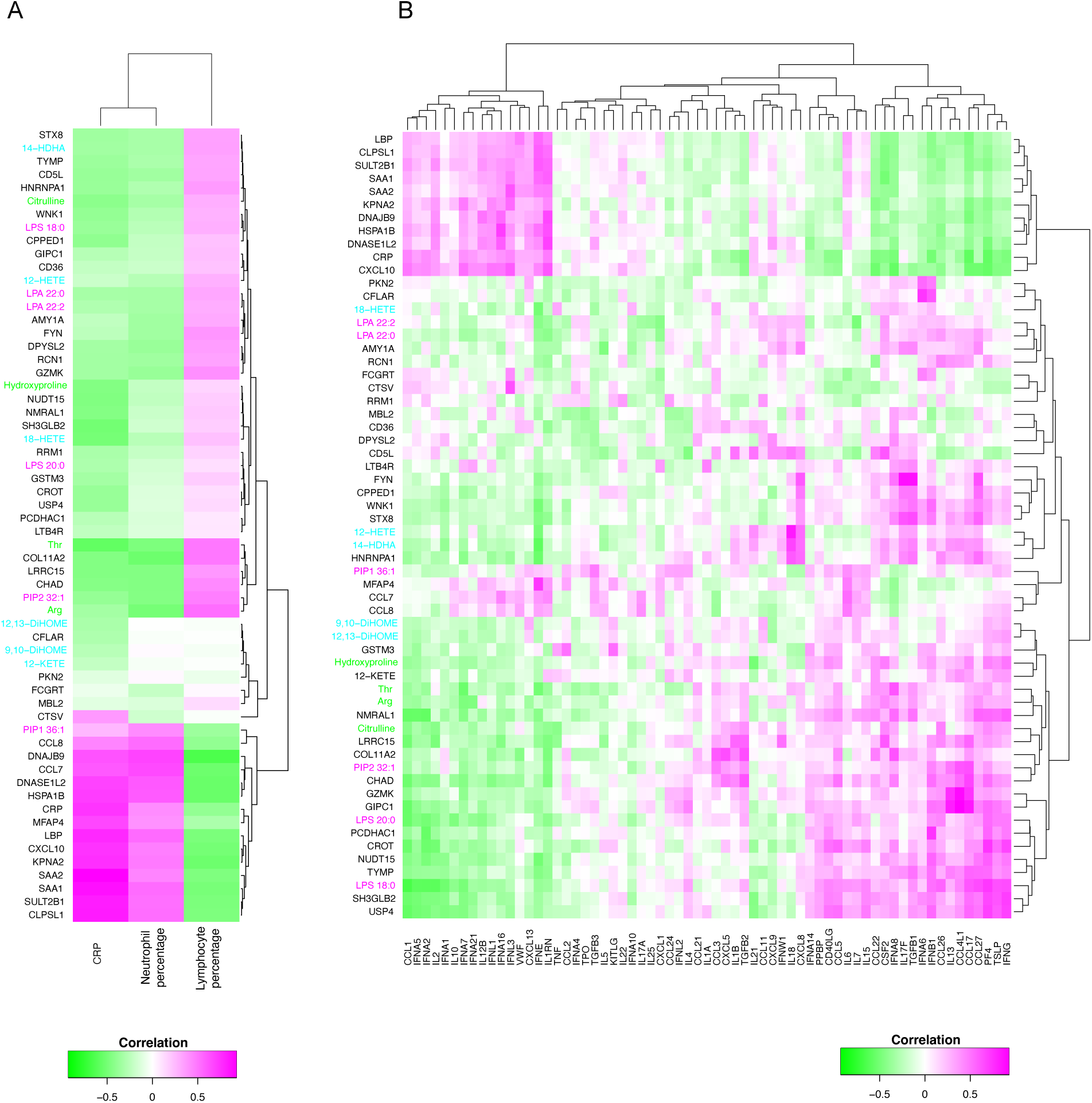
Associations between dysregulated molecules and clinical/immunological features. (A) Pearson correlation analysis between altered laboratory values and biomarker candidates. Proteins are labeled by their gene symbols. (B) Pearson correlation analysis between immunological markers and biomarker candidates. Lipids are indicated in red, lipid mediators in blue, aqueous metabolites in green, and proteins in black.

As discussed in the Introduction, impaired interferon (IFN) responses have been implicated in severe COVID-19. To examine this further, we assessed correlations between the altered molecules and immunological markers, including IFNs (types I, II, and III), cytokines linked to cytokine release syndrome (CRS), and additional immune markers [10,44]. Unsupervised clustering revealed distinct immunological response profiles (Fig. 5B; Table S7). Markers on the left side of the heatmap positively correlated with CRP and SAA1, indicating an inflammatory profile. Conversely, those on the right—including type I IFNs such as IFNA6 and IFNA8—showed negative correlations with inflammatory markers. This finding aligns with previous reports suggesting that deficient type I IFN responses may contribute to immune evasion and increased severity in COVID-19 [10,44].

## Discussion

In this study, we revealed multi-omics alterations in plasma between “stable” and “progressive” COVID-19 patients prior to disease exacerbation. The samples were collected in Japan between January and December 2020, during which the predominant circulating strains were B.1.1, B.1.1.284, and B.1.1.214 lineages [45,46]. Our analyses demonstrated a marked reduction in lipid mediators and polar metabolites in patients who subsequently developed severe disease. Proteomic profiling also clearly separated the groups via PLS-DA, and a distinct panel of dysregulated molecules showed strong correlations with clinical and immunological markers.

Several clinical laboratory parameters, such as neutrophil and lymphocyte counts, platelet count, and CRP, have been reported as severity-associated markers in COVID-19 [43,47]. Parameters such as AST and ALT are routinely used to assess liver damage and other organ dysfunction [48,49]. However, relatively few studies have addressed their value as predictive markers [50,51]. For example, CRP and the neutrophil-to-lymphocyte ratio (NLR) have been shown to predict ICU admission and mortality. Notably, our data supported the predictive value of CRP, neutrophil, and lymphocyte levels in daily medical practice—despite none of our “progressive” patients requiring mechanical ventilation.

Considering that lymphocyte ratio, but not counts, showed a statistically significant difference, the increase of neutrophil counts might be the essence of disease state in the early stage of COVID-19 progression.

Based on AUC values, we identified nine candidate biomarkers. Thr was significantly decreased in the “progressive” group, consistent with a previous report [21], and also showed the second strongest positive correlation with lymphocyte percentage (r = 0.49).

However, since the difference in Thr levels disappeared after admission, it may not be ideal as a clinical predictor of exacerbation.

CCL7 and CCL8, both monocyte chemoattractant proteins, were upregulated in severe cases—plausibly reflecting monocyte involvement in CRS. DNAJB9 and HSPA1B are stress-response proteins involved in preventing protein misfolding. MFAP4, an extracellular matrix protein of the fibrinogen-related superfamily [52], was also upregulated, in agreement with a prior study [53]. However, its levels also normalized after admission, limiting its utility as an early biomarker.

To our knowledge, CTSV, COL11A2, and DNASE1L2 have not previously been reported as severity-discriminating markers in COVID-19. Further studies are needed to validate their clinical relevance.

As discussed in the Introduction, evasion of innate immunity—particularly type I and III IFN responses—is a key driver of COVID-19 severity [10,44,54]. Viral proteins such as ORF3b and ORF9b inhibit type I IFN responses at multiple steps in the signaling cascade [55–57], and autoantibodies against IFN-α2 and IFN-ω have been reported in life-threatening cases [58]. In our dataset, IFN-α6, -α8, -α14, IFN-β1, and IFN-γ were negatively correlated with CRP and proinflammatory proteins like SAA1 and SAA2. Notably, IFN-α8 is induced by CpG motifs [59], and SARS-CoV-2 has the most pronounced CpG deficiency among known betacoronaviruses [60], potentially contributing to impaired IFN-α8 induction and immune evasion.

Type III interferons, such as IFN-λ1 and IFN-λ3, are essential for antiviral defense at epithelial surfaces (e.g., respiratory and gastrointestinal tracts). These IFNs showed positive correlations with CRP in our dataset, consistent with previous reports [61]. One study reported elevated IFN-λ3 levels in severe-to-critical patients. However, other reports have documented potentially harmful effects of IFNs; for example, a multicenter cohort study suggested that early administration of IFN-α2b reduced viral load, whereas late administration increased mortality [62]. One possible explanation is that IFN-α upregulates ACE2 expression, thereby enhancing viral entry into host cells [63,64]. Our findings may help refine therapeutic strategies involving IFNs.

This study has several limitations. It was conducted at a single center with a small sample size. The limited number of “progressive” patients is partly due to the overall low rate of progression to severe disease among hospitalized patients with mild-to-moderate illness. Additional limitations include an imbalanced sex distribution and the absence of healthy controls. Furthermore, AVL buffer was used for viral nucleic acid purification in the lipidomics workflow, which may have affected lipid extraction profiles.

In summary, we conducted a multi-omics analysis of plasma samples collected prior to clinical deterioration in COVID-19. Our results identified molecular signatures associated with disease progression and yielded candidate biomarkers with high diagnostic potential.

These findings provide new insights into the early pathophysiological processes of COVID-19 and may support future research on virus-induced acute exacerbation.

## Acknowledgements

We would like to express our gratitude to Human Metabolome Technologies, Inc. (HMT) and NEC Solution Innovators, Ltd.. for their cooperation in the analytical measurements. We are grateful for the constructive comments from K. Waku (Teikyo University), S. Morioka, N. Iwamoto, S. Saito (Japan Institute for Health Security). We would like to thank all the members in our laboratory (Japan Institute for Health Security) for their advice, discussions and technical support. Some parts of the English language in this manuscript were improved using a generative AI tool (ChatGPT), which was used solely for language editing and did not influence the scientific content.

## Data availability

Raw data and R scripts of this study are available in Zenodo at https://doi.org/10.5281/zenodo.17527930.

