## Supplemental information for "Molecular signature of COVID-19 prior to its exacerbation by multi-omics survey"

**Supplementary materials**

**Fig. S1. Molecular biomarker candidates after COVID-19 exacerbation**

(A-I) Bar plots after admission using plasma samples (means ± SEM) (***$p < 0.001$, *$p < 0.05$).

**Fig. S2. Clinical biomarker candidates after COVID-19 exacerbation**

(A-C) Bar plots after admission using plasma samples (means ± SEM) (**$p < 0.01$).

**Supplementary Table S1–S7: Analysis-derived results group**

Table S1 Dysregulated lipid molecules in the volcano plot

Table S2 Dysregulated lipid mediators in the volcano plot

Table S3 Dysregulated aqueous metabolites in the volcano plot

Table S4 Dysregulated proteins in the volcano plot

Table S5 Candidate clinical parameters

Table S6 Pearson correlations between signature molecules and significant clinical parameters

Table S7 Pearson correlations between signature molecules and reported immunological markers

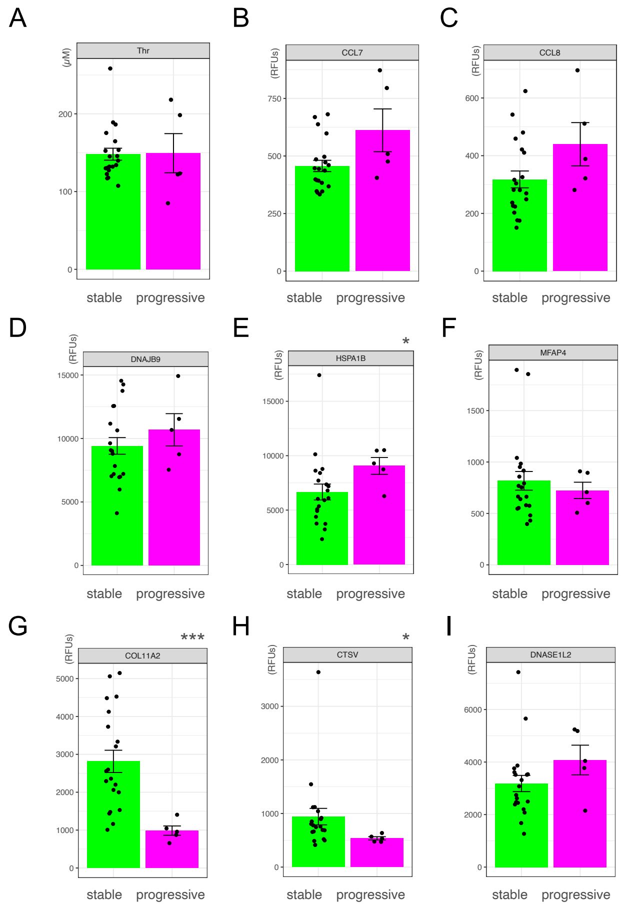


Fig S1. Molecular biomarker candidates after COVID-19 exacerbation

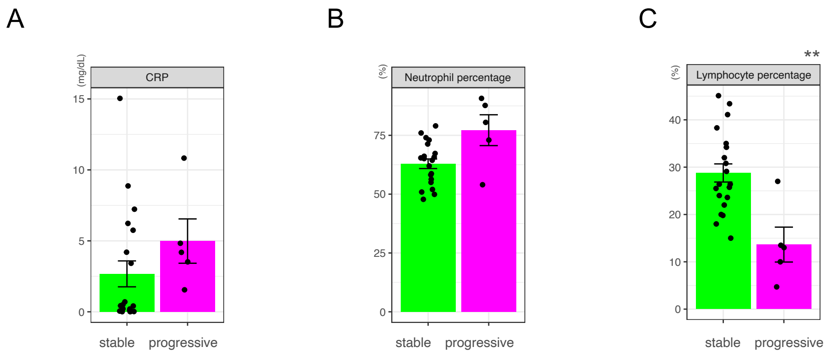


Fig S2. Clinical biomarker candidates after COVID-19 exacerbation
